# Traction force screening enabled by compliant PDMS elastomers

**DOI:** 10.1101/162206

**Authors:** Haruka Yoshie, Newsha Koushki, Rosa Kaviani, Kavitha Rajendran, Quynh Dang, Amjad Husain, Sean Yao, Chuck Li, John K Sullivan, Magali Saint-Geniez, Ramaswamy Krishnan, Allen J Ehrlicher

**Author notes:** contributed equally (Ramaswamy Krishnan, Allen J Ehrlicher).

## Abstract

Acto-myosin contractility is an essential element of many aspects of cellular biology, and manifests as traction forces that cells exert on their surroundings. The central role of these forces makes them a novel principal therapeutic target in diverse diseases. This requires accurate and higher capacity measurements of traction forces; however, existing methods are largely low throughput, limiting their utility in broader applications. To address this need, we employ Fourier-transform traction force microscopy in a parallelized 96-well format, which we refer to as contractile force screening (CFS).Critically, rather than the frequently employed hydrogel polyacrylamide (PAA), we fabricate these plates using polydimethylsiloxane (PDMS) rubber. Key to this approach is that the PDMS used is very compliant, with a lower-bound Young’s modulus of approximately 0.7 kPa. We subdivide these monolithic substrates spatially into biochemically independent wells, creating a uniform multiwell platform for traction force screening. We demonstrate the utility and versatility of this platform by quantifying the compound and dose-dependent contractility responses of human airway smooth muscle cells and retinal pigment epithelial cells.

## Introduction

Many adherent cells employ acto-myosin contractility to exert traction forces on their surroundings. These forces are an essential part of cellular deformation^[1]^, adhesion ^[2]^, spreading ^[3]^, and migration^[4]^, as well as growth ^[5]^, homeostasis ^[6]^, gene expression ^[7]^, and apoptosis ^[8]^. The significant role of traction force makes it a novel principal therapeutic target in diverse diseases, however, accurate measurements of traction forces are essential for this approach.

To quantify cell traction forces, researchers have employed a variety of techniques and tools. From the first wrinkling thin silicone sheets ^[9]^ to complex 3D multicellular contractility ^[10]^, a multitude of biomechanical methods have been developed collectively referred to as Traction Force Microscopy (TFM), as reviewed here ^[11]^. While these approaches have enabled the discovery of valuable mechano-biological connections, these methods are generally inherently slow and restricted to low-throughput implementation, limiting their utility as tools in broader pharmacological applications.

To address this need, we employ Fourier-transform traction force microscopy in a parallelized 96-well format, an approach we refer to as contractile force screening (CFS). Critically, rather than using the frequently employed hydrogel polyacrylamide (PAA), we fabricate these plates using polydimethylsiloxane (PDMS) rubber. Key to this approach is that the PDMS used is very compliant, with a lower-bound Young’s modulus of approximately 0.7kPa, unlike commonly used Sylgard 184 formulations. Like PAA, soft PDMS elastomers possess several material-favorable properties: their stiffness is tunable over a large physiological range (Fig 1), and they are non-toxic, non-degrading, and biologically inert. In addition to these aspects, this compliant PDMS has numerous advantages over PAA: 1) it is optically transparent with a refractive index of ~ 1.4 which is comparable to glass, 2) it is indefinitely stable after production without special storage considerations, 3) it is amenable to spin coating providing a simple means of creating a uniform and flat surface, 4) it is a uniformly non-porous surface, unlike PAA whose porosity can vary strongly with cross-linking concentration ^[12]^.Critically, the monolithic impermeable nature of these silicone substrates makes them easy and ideal to subdivide spatially into biochemically independent wells, creating a uniform multi-well platform for traction force microscopy. Taken together, our compliant PDMS presents numerous advantages in becoming a new standard in soft substrates for TFM; these advantages are particularly important for a standardized higher-throughput technology and will enable widespread adoption of CFS from our previous approach using PAA ^[13]^

**Figure 1:**
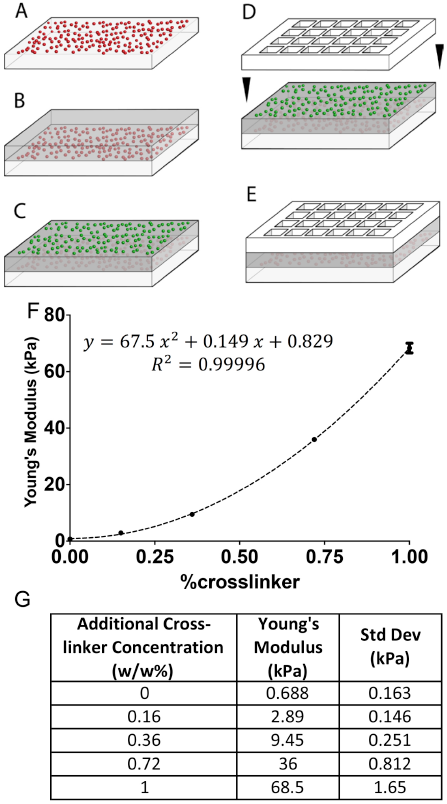
Fabrication of multi-well substrates and mechanical characterization of tunable PDMS elastomer. Multi-well PDMS devices for TFM are fabricated by A) optionally coating a layer of fluorescent microspheres (for de-drifting images) on the custom glass slide, B) spin coating a ~100μm thick layer of compliant PDMS, C) spin coating a ~1μm thick layer of compliant PDMS mixed with custom-synthesized beads with different fluorescence emission for measuring cell induced deformation of the PDMS, D) bonding a multiwell partitioning strip to the top, and E) ligating and culturing cells. F) PDMS moduli as determined by nanoindentation (n=14-16 measurements per data point). G) quadratic fit of Young’s moduli of PDMS as a function of additional cross-linker.

## Methods

### Cell culture

Primary human airway smooth muscle (ASM) cells were obtained from the Gift of Hope Organ and Tissue Donor Network. These cells have been well-characterized previously, e.g. ^[14]^. All measurements were performed using cells at passage 5-8 from two non-asthmatic donors. ARPE-19 (retinal pigment epithelium) cells were obtained from American Type Culture Collection. All culture media formulations are provided in the supplemental material.

### Preparation of silicone substrates in custom 96-well plates

We fabricate our multiwell TFM dishes by applying very compliant and tunable modulus PDMS onto custom cut glass slides and then partitioning the wells with a plastic subdivider. In brief, very compliant commercial PDMS (NuSil® 8100, NuSil Silicone Technologies, Carpinteria, CA) is mixed with Sylgard 184 crosslinking agent to make a tunable (E= 0.68 to 68 kPa) substrate, which is impregnated with a ~1μm thick layer of fiduciary particles to reveal cell-induced deformations. This approach differs from existing PDMS TFM strategies, as this substrate is comparably compliant to polyacrylamide and linearly elastic, yet not a hydrogel. Full details of plate preparation are provided in the supplemental materials.

### Measurements of cell traction forces

The 96-well plate was mounted within a heated chamber (37°C) upon an automated computer-controlled motorized stage and imaged at 10x magnification using a monochrome camera (Leica DFC365 FX) affixed to an inverted microscope (DMI 6000B, Leica Inc., Germany). We acquired fluorescent images of microspheres embedded in the elastic substrate immediately underneath the cells at (i) *baseline* with no treatment, (ii) after *treatment*, and (iii) after cell detachment with trypsin (*reference null-force image*). By comparing the fluorescent images at reference with the corresponding images at baseline and after treatment, we obtain a time series of bead-displacement, and hence substrate deformation fields (resolution = ~15μm). Using the measured substrate deformation, the pre-defined substrate modulus, and thickness, traction force maps and the root-mean squared value were calculated on a well-by-well basis using the approach of Fourier-transform traction cytometry ^[15]^ modified to the case of cell monolayers ^[16]^.

### Drugs

Histamine, Isoproterenol, Salbutamol, Salmeterol, Formoterol, Thrombin, and H_2_0_2_ were purchased from Sigma-Aldrich. Y27632 was purchased from EMD Millipore. Human VEGF-A^165^ and Bevacizumab were purchased from R&D systems and Genentech, respectively.

### Statistics

Statistical comparisons were performed using the non-parametric Wilcoxon matched-pairs signed rank test. Differences were considered significant when p<0.05.

## Results and Discussion

CFS entails quantifying the cell-generated forces by measuring fluorescent bead positions in each well of the 96-well plate: 1) without cells, 2) with cells adhered at baseline contractility, and 3) after treatment with the compound(s) of interest. For example, for a representative well of a 96-well plate (Fig. 2A), shown are ASM traction force maps and the root-mean squared value (inset) at baseline (Fig, 2D, 73Pa), following treatment with the contractant, the H1 agonist, histamine (Fig. 2E, 88Pa), and after additional treatment with the relaxant, the β2 adrenergic receptor agonist, isoproterenol (Fig. 2F, 41Pa).

**Figure 2:**
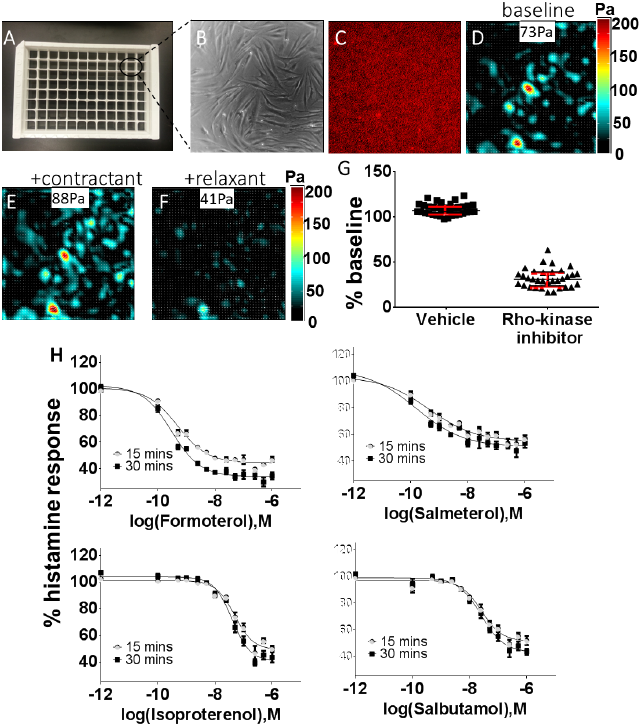
CFS using soft elastomeric substrates recapitulates known ASM pharmacological responses. Human ASM cells were cultured to confluence upon 9.45kPa stiff collagen-coated 96-well silicone substrates. A-D) For a representative well of a 96-well plate, shown are images of cells, fluorescent beads, traction force maps and average magnitude (inset) at baseline. E-F) For the same well, shown are traction force maps and average magnitude (inset) with the contractant compound, histamine (10μM, 30 minutes), and after additional treatment with the relaxant compound, Isoproterenol (0.5μM, 30 minutes). G) Over the 96-well plate, the force measurements are statistically different (*p*<0.05) between positive and negative controls, as ascertained by an unpaired t-test. H) Force measurements confirmed known differences in potency amongst a panel of functionally diverse ASM relaxation compounds (Formoterol > Salmeterol > Salbutamol > Isoproterenol). Plotted are the mean±SEM calculated from 3-8 wells per dose per ASM relaxation compound. Data were pooled from 2-4 96 well plates tested on different days but under identical experimental conditions.

First, we tested the suitability of our approach for higher-capacity measurements. Individual wells of a single 96-well plate were either assigned to a *positive* or *negative control* group. In the positive control group, cells were pre-stimulated with 10 μM histamine to induce maximal contractility followed by post-stimulation with the relaxant, 10μM Y27632 for 30 minutes. In the negative control group, cells were pre-stimulated with vehicle (PBS) followed by post-stimulation with vehicle (PBS) for 30 minutes. In both groups, traction force measured post-stimulation was normalized to the corresponding pre-stimulation value on a well-by-well basis. From these measurements of normalized changes, we determined that the groups were statistically different (*p*<0.05), as ascertained by an unpaired student t-test.

Next, we verified the utility of our approach in pharmacology by examining traction force changes induced by a diverse set of well-known and clinically relevant airway smooth muscle (ASM) relaxation compounds ^[17]^. Each compound was evaluated in a 10-point dose response manner, across adjacent rows of the 96 well-plate. Data were pooled from multiple plates and reported as a percentage of histamine response. The extent of ASM relaxation confirmed the known differences in potency of the β2 adrenergic receptor agonists (Salmeterol > Formoterol > Salbutamol > Isoproterenol) ^[17]^, and the full agonist, Formoterol, provided a greater scope of relaxation than the partial agonist, Salmeterol, as expected ^[18]^ (Fig. 2H, Supplementary Table 1). Notably, as supported by negligible standard errors and the small coefficients of variation, the data were highly reproducible.

Here we have focused on ASM response; yet this approach is applicable in pharmacology to any adherent contractile cell type and is therefore expected to be of broad utility. In ASM this need is particularly exigent, as current efforts to screen new ASM relaxation drugs employ indirect assay methods that are poorly predictive of functional response. Commonplace examples include the dissociation of intracellular calcium regulation from the effects of bradykinin, bitter tastants ^[19]^, and proton-sensing receptor ligands ^[20]^ on ASM contraction, a similar dissociation of cAMP regulation from bronchorelaxant effect (pro-contractile receptor antagonists, and again bitter tastants), and the limited predictive utility of membrane potential for almost all drugs whether they target receptors or other contractile effectors or signaling elements. A more relevant screen that directly quantifies the target output of ASM relaxation, as does CFS, is required to efficiently test the pending generations of ASM relaxation drugs. To this end, CFS fills an important methodological void in ASM biology, and more generally, in measuring contractile response for a broad range of cell types and pathologies.

To demonstrate the versatility of CFS, we examined a key pathogenic mechanism common to many ocular pathologies – dysfunction of the retinal pigmented epithelium (RPE) ^[21]^. We discovered that the RPE barrier-disruptive agent, thrombin ^[22]^, the pro-angiogenic cytokine, VEGF-A ^[23]^, and the oxidative stressor, H_2_O_2_ ^[24]^, each caused an increase in RPE traction forces (Fig. 3). Conversely, the Rho Kinase inhibitor, Y27632, or the VEGF-A inhibitor, Bevacizumab, ablated these forces. Taken together, these data reveal a novel role for traction force enhancement in RPE dysfunction and advocate for new discovery efforts targeted at reducing these forces. This might be especially pertinent to offset RPE dysfunction in the commonly occurring dry form of macular degeneration ^[25]^, wherein no therapeutic intervention currently exists.

**Figure 3:**
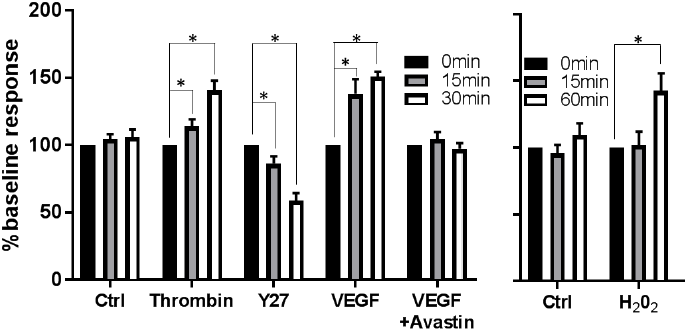
Mediators of retinal epithelial dysfunction promote cell traction forces. Human ARPE-19 cells were cultured to confluence upon 9.45kPa stiff collagen-coated 96-well silicone substrates, and cell-contractile forces were measured at baseline (0 min) and after treatment (15, 30/60 min). While thrombin (1unit/ml) VEGF (100ng/ml), and H_2_0_2_ (100nM) enhanced baseline forces in a time-dependent manner, the rho-kinase inhibitor, Y27632 (10μM) ablated them. Bevacizumab (0.05mg/ml) prevented the VEGF-induced force enhancement. Quantitative average changes were pooled from 8-24 well per time point per treatment. ^*^ indicates significant difference compared to baseline.

## Conclusion

We have demonstrated utility for a 96-well silicone-based substrate for CFS. This approach is advantageous over CFS using PAA ^[13]^ as the material itself is more robust and uniform, and the fabrication of multiwell-silicone substrates eliminates time-consuming production steps, utilizes standard micro-fabrication procedures, and obviates the need for surface-bound fluorescent beads by embedding them as a monolayer by spin-coating. Moreover, silicone elastomers possess many material-favorable properties over PAA. Specifically, they are predominantly elastic in the stiffness range that encompasses most physiological microenvironments, are non-porous and impermeable, thus obviating common concerns associated with PAA ^[12]^, and possess superior optical properties.

Mechanical malfunction appears be an integral component of many diverse diseases including asthma, ocular pathologies, acute lung injury, bladder dysfunction, vascular diseases, fibrosis, and cancer, wherein cell contractile forces play a pivotal role. CFS using elastic silicone substrates is expected to enable mechanistic studies for both quantitatively describing the aberrant contractile forces, as well to mechanically rectify the responses by directly evaluating potential therapeutic compounds.

## Acknowledgements

Cynthia Crespo and Erin Hannen for technical assistance. AJE acknowledges NSERC RGPIN/05843-2014 & EQPEQ/472339-2015, CIHR Grant # 143327, and CCS Grant #703930. RK acknowledges

NIH R21HL123522 and Amgen Inc. HY was supported by FRQS. MSG acknowledges NIH DP2-OD006649.

